# ASK1/p38-mediated NLRP3 inflammasome signaling pathway contributed to aberrant retinal angiogenesis in diabetic retinopathy

**DOI:** 10.1101/763102

**Authors:** Wenjun Zou, Zhengwei Zhang, Shasha Luo, Libo Cheng, Xiaoli Huang, Nannan Ding, Jinjin Yu, Ying Pan, Zhifeng Wu

## Abstract

Diabetic retinopathy is the leading cause of blindness in the working-age population in many countries. Despite the available treatments, some patients present late in the course of the disease when treatment is more difficult. Hence, it is crucial that the new targets are found and utilized in clinical therapy of diabetic retinopathy. In this study, we constructed the DR animal model and the high model in HRMEC cell to investigate the relationship between ASK1/p38 and NLRP3 in DR. The results showed that DR could cause the inflammatory response and microvascular proliferation. NLRP3 contributed to DR-mediated inflammatory development and progression, which promoted the inflammatory related cytokine expression. Meanwhile, it could promote the tube formation of retinal microvascular endothelial and angiogenesis. Moreover, further research showed that NLRP3 mediated aberrant retinal angiogenesis in diabetic retinopathy was regulated by ASK1 and p38. It suggested that ASK1/p38 could become a new target in DR treatment.

## Introduction

Diabetic retinopathy (DR) is a common and specific microvascular complication of diabetes, and it remains the leading cause of preventable blindness in working-aged people [1,2]. It is associated with diabetes and increases the risk of life-threating systemic vascular complications, which include stroke, coronary heart disease, and heart failure [3,4]. The main reason for the irreversible visual induced by diabetic retinopathy is retinal neovascularization [5]. DR is usually related with some signaling pathway disorder and the unusual functional molecular expression [6]. Hence, potential molecular mechanism of retinal neovascularization is of great importance and it is conducive to developing effective diagnosis and therapy technologies for DR patients.

Some reports show that DR can cause the activation of polyol pathway and hexamine pathway, accumulation of advanced glycation end products, inflammation and protein kinase C [5,7,8]. More and more evidence show that inflammatory response plays an important role in the pathogenesis of DR [7]. Moreover, the high expression levels of pro-inflammatory cytokines are examined in retinas from animals with diabetes [5]. Therefore, the inhibition of inflammatory signaling can become a treatment for DR therapy.

The inflammasome is a multiprotein scaffolding complex which includes a member of NOD-like receptor family, pyrin domain containing family member (NLRP), procaspase 1 and apoptosis associated speck-like protein containing CARD [9]. To data, NLRP3 inflammasome can leads to secretion of pro-inflammatory cytokine IL-1β through recognizing danger signals and apoptosis associated speck-like protein to activate caspase-1[9]. Related reports illustrate that NLRP3 activation play a crucial role in metabolic disease, such as type 2 diabetes [10]. Moreover, clinical samples in various stages of diabetic patients show that expression of the NLRP3 and related inflammatory protein are increased in vitreous sample, which have the largest response in patients with proliferative DR [11]. Thus, it appears that the NLRP3 inflammasome may be involved in retinal disease.

Apoptosis signal-regulating kinase 1 (ASK1), a member of the mitogen-activated protein kinase kinase kinase (MAP3K) family, is an important stress responsive protein kinase which play crucial role in the initiation of many diseases including neurodegenerative, cardiovascular, inflammatory, autoimmunity, and metabolic disorders [12,13]. The MAPK member, p38, is a serine/threonine protein kinase, which responds to several cellular processes and external stress signaling, such as cell differentiation, cell proliferation, inflammation regulation and cells death [14,15]. ASK1 is the most studied family member and is an upstream kinase of the JNK and p38 pathways [16,17]. Previous studies shows that ASK1 is activated in response to a variety of stress-related stimuli *via* distinct mechanisms and activates MKK4 and MKK3, which in turn activate JNK and p38 [18]. Related studies reports that ASK1/2 signaling complex contributes to pyroptotic cell death by regulating the NLRP3 inflammasome [19]. Although ASK1/p38 play a substantial role in inflammatory, and it contribution in DR pathogenesis has not been described.

In this study, we constructed the DR animal model and the high model in HRMEC cell to investigate the relationship between ASK1/p38 and NLRP3 in DR. We found that ASK1/p38 could mediate NLRP3 inflammasome signaling pathway contributed to aberrant retinal angiogenesis in DR. It suggested that ASK1/p38 could become a new target in DR treatment.

## Material and methods

### Cells culture and stimulation

The Primary Human Retinal Microvascular Endothelial cell (HRMECs) was purchased from Cell systems (Kirkland, WA, USA), and routinely cultured in M199 medium (Millipore, Temecula, CA) supplemented with100 units of penicillin and 100 μg of streptomycin per milliliter of medium. All cells (passages 5 – 12) were cultured in grade plastic-ware and maintained in an atmosphere of 5% CO_2_ at 37°C.

For high glucose cell models, HRMECs were cultured in conditioned medium with 5 mM (serving as the normal glucose (NG) group) or 30 mM (HG group) D-glucose (Sigma, Darmstadt, Germany) and incubated at 37°C with 5% CO_2;_ then, HG group was treated with or without 1 μM NLRP3 inhibitor CY-09 (MedChemExpress, New Jersey, USA), 1 μM ASK1 inhibitor GS-444217 (MedChemExpress, New Jersey, USA) or 10μM p38 inhibitor SB203580 (MedChemExpress, New Jersey, USA) for 24 h, respectively. Each inhibitor was dissolved in dimethyl sulfoxide (DMSO) to a 50 mM concentration for use as stock solutions that were diluted to the required concentrations for in vitro studies.

### The constructive and treatment of DR model

Male C57/BL/J (10 weeks old, Yangzhou, Jiangsu, China) were obtained and housed under standard conditions at Animal Research Center of China Pharmaceutical University. All animal experiments were performed according to protocols approved by the Ethics Committee of China Pharmaceutical University. Streptozotocin (STZ)-induced hyperglycemic mice were utilized as type I diabetic-like model associated with retinopathy [20]. Male C57/BL/J mice were received 1 times/day constitutive intraperitoneal injections of 50 mg/kg STZ in a citric buffer (pH 4.5) for 5 day. After last injection, 4 hour fasting blood glucose was determined, which the fasting blood glucose were included within 15.0 to 20.0 nmol/l. The control group mice were injected only citric buffer. Hyperglycemic (HG) mice were randomized divided into 5 groups: model and NLRP3 inhibitor CY-09 treatment groups, ASK1 inhibitor GS-444217 treatment groups and p38 inhibitor SB203580 treatment groups. NLRP3 inhibitor CY-09 was administrated by oral administration at a dose of 2.5 mg/kg once a day for 6 weeks in C57BL/6J mice. ASK1 inhibitor GS-444217 was administrated by oral administration at a dose of 10 mg/kg once a day for 6 weeks in C57BL/6J mice. The p38 inhibitor SB203580 was administrated by oral administration at a dose of 15 mg/kg once a day for 6 weeks in C57BL/6J mice. Normal and model groups of mice received the same volume of water. All inhibitors were first dissolved in a small quantity of DMSO, and then evenly mixed with 0.5% CMC-Na solution. The final concentration of DMSO was 5%. Treatment lasted 6 weeks. Eyeballs were collected from sacrificed mice. Some retina was enucleated and placed on 4% paraformaldehyde overnight for immunofluorescence. Another retina was reversed in −80°C for qRT-PCR or Western blotting experiment.

### Quantitative real-time PCR analysis

Total RNA of HRMECs and tissue samples were extracted using TRIzol and was transcript into cDNA with a Reverse Transcription Kit (Bio-Red). Then, qRT - PCR was then performed on a MyiQ single-color RT - PCR detection system with SYBR Green Supermix. Primer sequences for each gene were as follow: IL-6, forward: 5’- CTT CAC AAG TCG GAG GCT TAA T-3’, reverse: 5’- GCA TCA TCG CTG TTC ATA CAA TC-3’; TNF-α, forward: 5’- GCC TCA GCC TCT TCT CAT TC-3’, reverse: 5’- GGG AAC TTC TCC TCC TTG TTG-3’; IL-1β, forward: 5’- TGA CCC ATG TGA GCT GAA AG-3’, reverse: 5’ - CGT TGC TTG TCT CTC CTT GTA-3’; GAPDH, forward: 5’- GGG AAA CCC ATC ACC ATC TT-3’, reverse: 5’- CCA GTA GAC TCC ACG ACA TAC T-3’. The relative mRNA expression was normalized to that of GAPDH and was determined using the comparative Cq method (2-ΔΔCq) [21].

### Western blot analysis

For western blot analysis, the protein of HRMECs and tissue samples were lysed in RIPA buffer containing protease inhibitor cocktail (Generay, Shanghai, China). NLRP3, AKS1, p38, IL-6, TNF-α, VEGF, IL-1β and GAPDH were detected by western blot analysis with anti- NLRP3, anti- AKS1, anti- p38, anti- IL-6, anti- TNF-α, anti- VEGF, anti- IL-1β and anti- GAPDH (Cambridge, MA, USA). The blots were detected using Bio-Imaging System and Quality One 1-D analysis software (Bio-Rad, Richmond, CA, USA).

### Immunofluorescence staining

The cry sections from mouse retina tissues were fixed with ice-cold acetone for 20 min. The slides were blocked with 5% BSA in PBS for 1 h and subjected to incubation at 4°C overnight with the following primary Ab mixtures: biotin–anti-mouse CD31 (1:100) or biotin–anti-mouse IB4 (1:100). Slides were washed and then incubated with streptavidin-Alexa Fluor 488 conjugate (1:200) or streptavidin–Alexa Fluor 594 conjugate (1:200) for 90 min. The slides were costained with DAPI and mounted with fluoro-gel (Electron Microscopy Science). Confocal images were acquired by Leica TCS SP5 confocal microscope system (Leica Microsystems, Germany) and quantitated by AxioVision 4.6.3.0 (Carl Zeiss AG, Germany).

### The experiment of retinal microvascular endothelial cell tube formation

The 100 μl Matrigel per well was evenly spread on the bottom of 24-well plate and allowed to solidify at 37°C with 5% CO_2_ for 2 h. Different treated HRMECs, including NG group, HG group and HG treated with inhibitor group, were digested and centrifuged at 1000 g; the supernatant was removed and the cells were suspended in complete medium; the cells were separately seeded in 24-well plates at 3 × 10^5^ cells per well and cultured at 37°C with 5% CO_2_ for 8 h. The tube formation of retinal microvascular endothelial cell was photographed and analyzed by ImageJ software Angiogenesis Analyzer Plugin.

### Measurements of VEGF

The retinal tissue from control group, HG group and HG treated with inhibitor group were added in PBS and homogenized; the homogenate was centrifuged at 12000 g at 4°C for 10 min. Then, the supernatant was tested for VEGF by mouse VEGF ELISA kit (R&D Systems, MN, USA). For cells experiment, the cells cultured supernatant in different treated groups was harvested and tested by mouse VEGF ELISA kit.

### Statistical analysis

The data were statistical analysis by Student’s t-test using Graph pad prism 4.0 (Graph pad Software, La Jolla, CA). P< 0.05 was considered statistically significant.

## Results

### The inflammatory response and microvascular proliferation were caused by DR

Related reports show that DR can activate some signaling pathway and cause inflammation response [7]. Hence, the Streptozotocin (STZ)-induced hyperglycemic mice were utilized as type I diabetic-like model associated with retinopathy and constructed. The inflammation cytokines related with DR were measured; the results showed that the mRNA expression levels of IL-6, TNF-α and IL-1β were enhanced in retina of DR groups compared with control group (Figure 1A). Then, the formation of microvascular in retina was analyzed, and the expression of vascular marker CD31 was increased in DR group by confocal detection (Figure 1B). Meanwhile, the VEGF secretion levels in DR group was increased compared with control group (Figure 1C). All of these data suggested that DR could cause inflammatory response and microvascular proliferation in retina.

**Figure 1.**
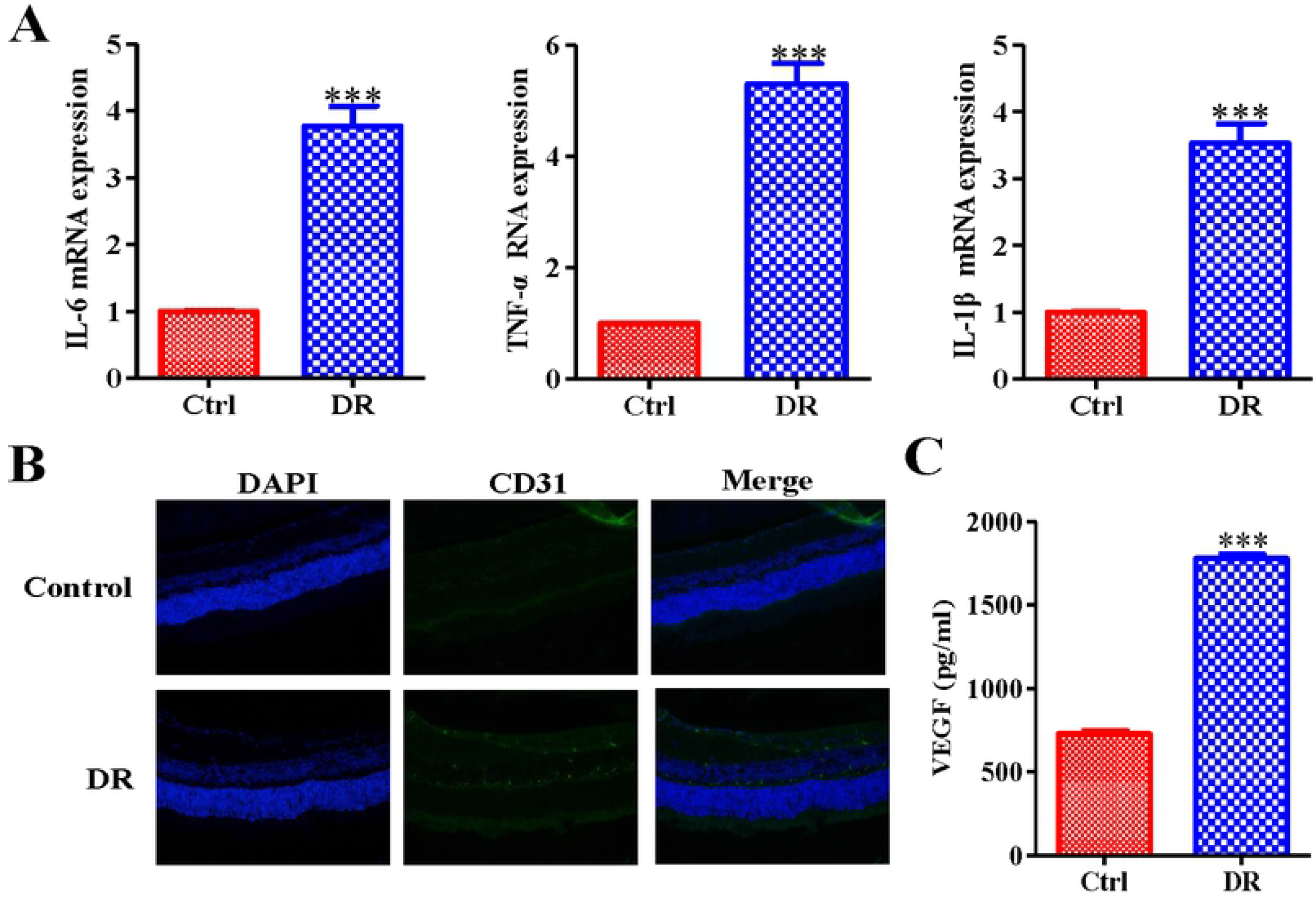
DR could cause the inflammatory response and microvascular proliferation. (A) The inflammatory related cytokine IL-6, TNF-α and IL-1β mRNA expression levels were enhanced in DR group. (B) The expression of vascular marker CD31 was increased in DR group by confocal detection. Meanwhile, (C) the VEGF secretion levels in DR group was increased compared with control group. Values are expressed as the mean ± SD (n = 6, *P < 0.05, **P < 0.01 and ***P < 0.001 compared with the control).

### NLRP3-mediated tissue inflammatory response promoted microvascular proliferation in retina

Clinical samples in various stages of diabetic patients show that expression of the NLRP3 and related inflammatory protein are increased in vitreous sample, which have the largest response in patients with proliferative DR [10]. Hence, in this study, we investigated the role of NLRP3 in microvascular proliferation in retina. The results showed that the protein expression of NLRP3 was up-regulated in DR group (Figure 2A). Then, the DR group was treated with NLRP3 inhibitor; after inhibitor treatment, the inflammation cytokines IL-6, TNF-α and IL-1β mRNA expression levels were decreased comparing with DR group without treated with inhibitor (Figure 2B). Meanwhile, CD31 expression level was reduced in inhibitor treatment group (Figure 2C). Moreover, the VEGF secretion level was reduced after administrated with NLRP3 inhibitor (Figure 2D). In order to further investigate the role of NLRP3, the high glucose cell models in HRMECs was constructed and treated with or without NLRP3 inhibitor. The results showed that NLRP3 protein expression levels in HG HRMECs was up-regulated, and inhibited after treated inhibitor (Figure 2E). In HG HRMECs groups, IL-6, TNF-α and IL-1β mRNA expression levels were enhanced comparing with control, and inhibited by NLRP3 inhibitor (Figure 2F). The result of VEGF secretion levels in cells was similar with DR animal model (Figure 2G). Taken together, this data illustrated that NLRP3-mediated tissue inflammatory response promoted microvascular proliferation in retina.

**Figure 2.**
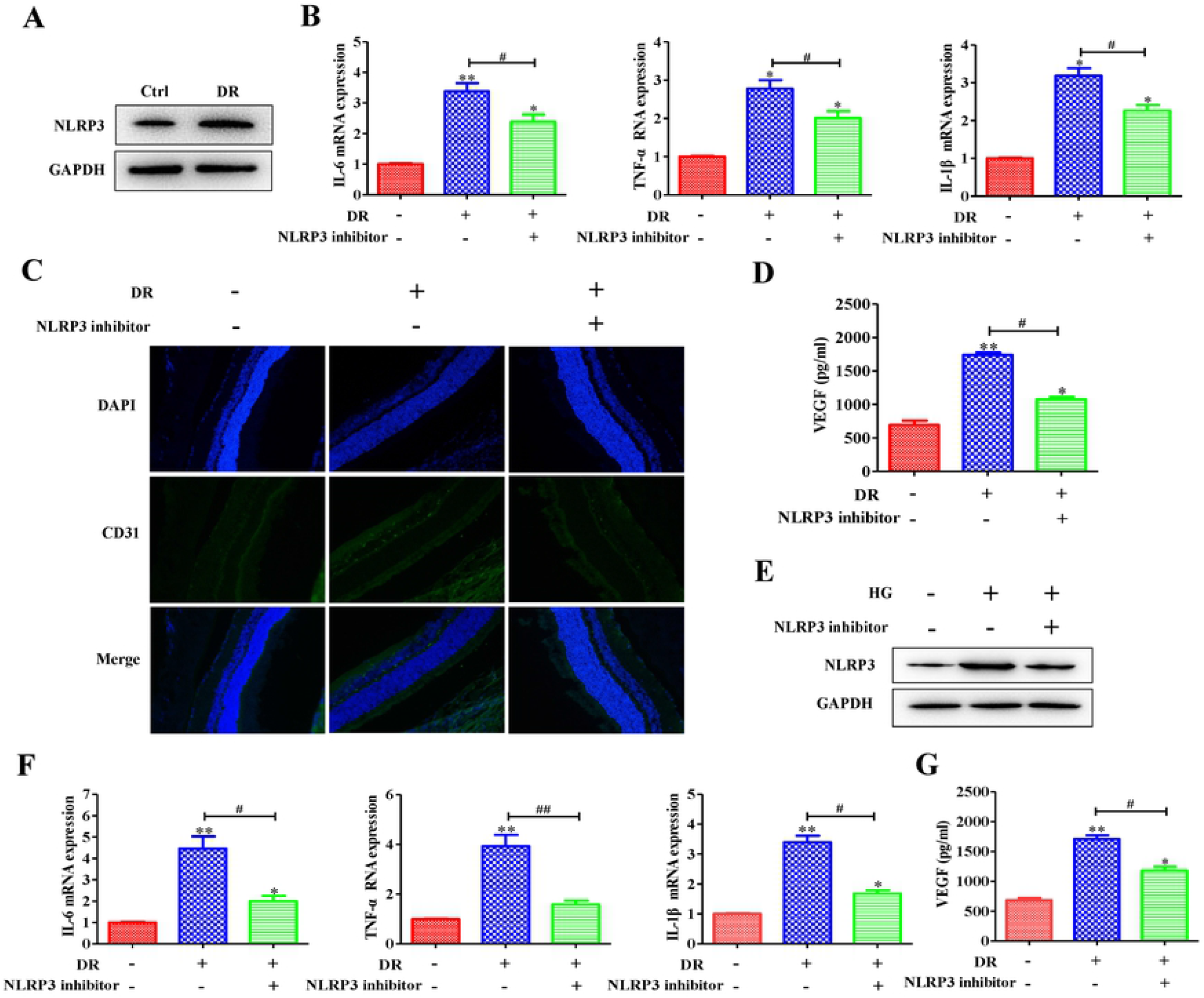
NLRP3-mediated tissue inflammatory response promoted microvascular proliferation in retina. (A) The protein expression level of NLRP3 was enhanced in DR groups; moreover, (B) blocking NLRP3, the mRNA expression of IL-6, TNF-α and IL-1β was down-regulated; meanwhile, (C) the vascular marker CD31 expression level and (D) VEGF secretion level were decreased through inhibiting NLRP3. In addition, in HRMEC cell model, (E) high glucose enhanced the protein expression of NLRP3, which was inhibited by NLPR3 inhibitor. (F) The inflammatory related cytokine IL-6, TNF-α and IL-1β was enhanced in HG-induced HRMEC cell model and was inhibited through blocking NLRP3. (G) The VEGF secretion level was enhanced, and was decreased after inhibiting NLRP3. Data are presented as the mean ± standard deviation from triplicate wells. *P < 0.05 and **P < 0.01 compared with the control. #P < 0.05 and ^##^P < 0.01 compared with the relative DR animal model group or HG-induced HRMEC cell group.

### The tube formation of retinal microvascular endothelial was inhibited through blocking NLRP3

Above data showed that NLRP3 was related with inflammatory response and promoted vascular marker CD31 and VEGF expression. Hence, we analyzed the tube formation of retinal microvascular endothelial in retina. Figure 3A showed that the lumen formation was obviously enhanced in HG HRMECs and was inhibited by blocking NLRP3. Then, the tube meshes, nodes and tube length was counted, and the results exhibited that HRMECs in high glucose could form more meshes and nudes than control; meanwhile, the tube total length in HG group was higher than control group (Figure 3B-C). However, after blocking NLRP3, the tube meshes, nudes and tube length was reduced comparing with HG group (Figure 3B-C). These data suggested that the tube formation of retinal microvascular endothelial was inhibited through blocking NLRP3.

**Figure 3.**
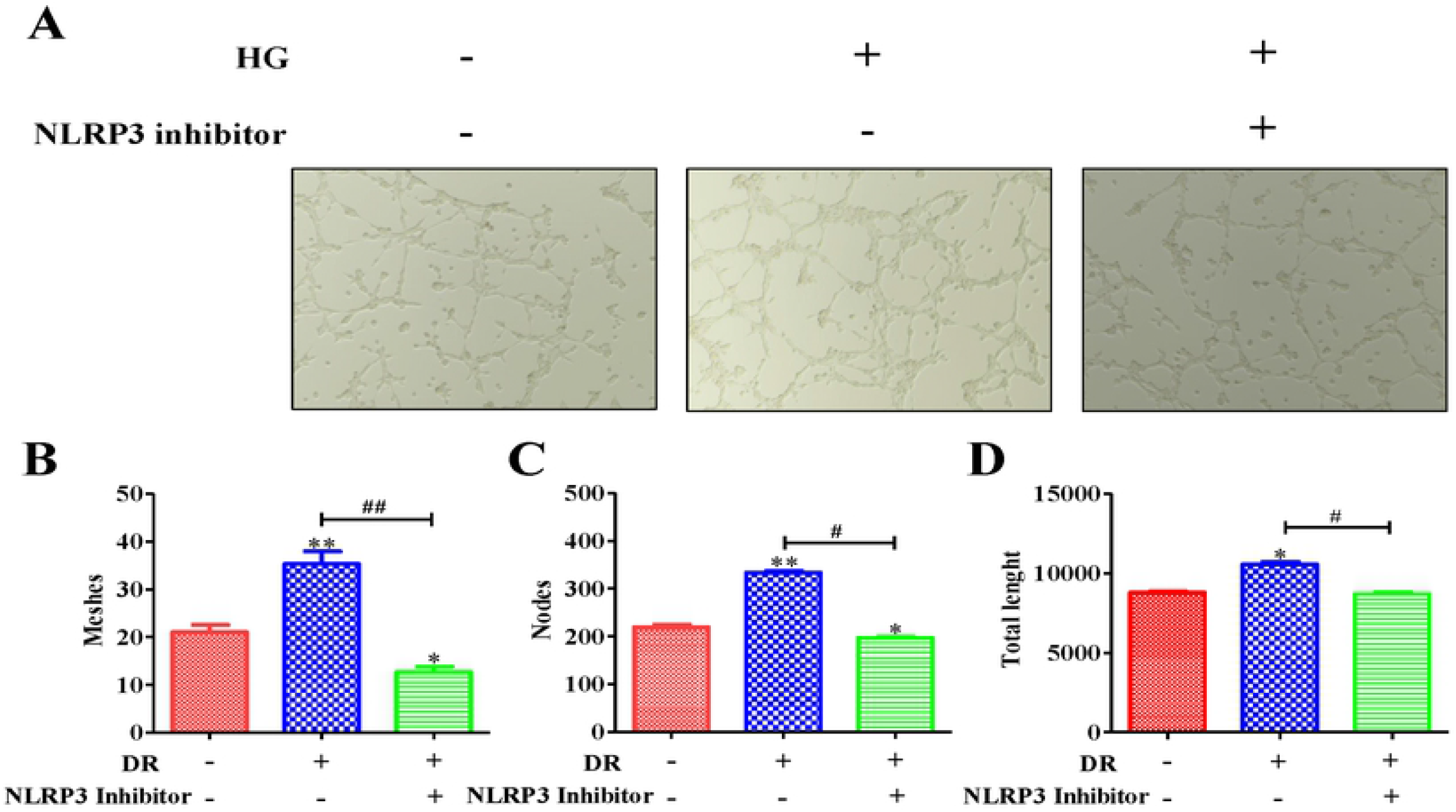
The tube formation of retinal microvascular endothelial was inhibited through blocking NLRP3. (A) The tube formation was obviously enhanced in HG HRMECs and was inhibited by blocking NLRP3. Then, the (B) tube meshes, (C) nodes and (D) tube length was counted, and the results exhibited that HRMECs in high glucose could form more meshes and nudes than control; meanwhile, the tube total length in HG group was higher than control group. However, after blocking NLRP3, the tube meshes, nudes and tube length was reduced comparing with HG group. *P < 0.05 and **P < 0.01 compared with the control. #P < 0.05 and ^##^P < 0.01 compared with the relative HG-induced HRMEC cell group.

### The expression level of inflammatory some NLRP3 was up-regulated through activating ASK1/p38 signal axis

More and more data show that NLRP3-induced inflammatory play an important part in DR [9]. However, the potential mechanism of NLRP3 participating in inflammatory in DR still unknown. So, we utilized the DR model and high glucose HRMECs model to investigate. The results showed that the protein expression levels of ASK1 and p38 were up-regulated in DR and high glucose HRMECs (Figure 4A-B). Then, in order to investigate whether ASK1 and p38 participating in NLRP3-mediated inflammatory response. We utilized ASK1 and p38 inhibitor to block the related protein expression; the data exhibited that the protein expression levels of NLRP3, inflammatory cytokine (IL-6, TNF-α, IL-1β) and VEGF was down-regulated after blocking ASK1 and p38 (Figure 4C-D). All of this data suggested that NLRP3-mediated tissue inflammatory signal response was enhanced through activating ASK1/p38 signal axis.

**Figure 4.**
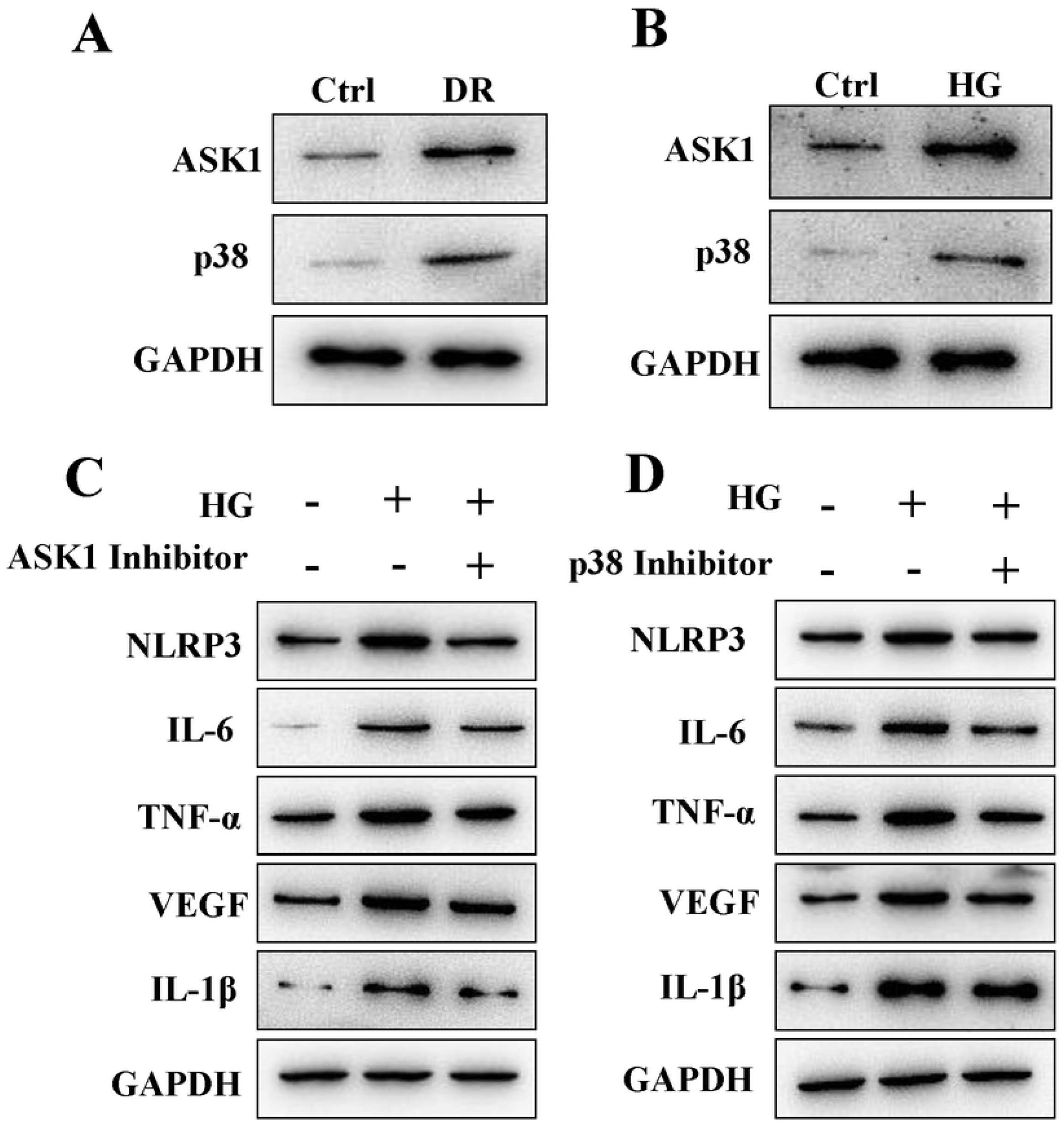
The expression level of inflammatory some NLRP3 was up-regulated through activating ASK1/p38 signal axis. (A) The protein expression levels of AKS1 and p38 were up-regulated in DR animal group. (B)The result of HG-induced HRMEC group was similar with DR animal group. Moreover, (C) the NLPR3, IL-6, TNF-α, IL-1β and VEGF protein expression levels were inhibited by (C) NLPR3 inhibitor and (D) p38 inhibitor.

### The tube formation of retinal microvascular endothelial was inhibited through blocking ASK1/p38 signal axis

NLRP3 play an important part in tube formation of retinal microvascular endothelial. Meanwhile, NLRP3-mediated tissue inflammatory signal response was enhanced through activating ASK1/p38 signal axis. Hence, we hypothesized whether ASK1/p38 signal axis could mediate tube formation of retinal microvascular endothelial. Figure 5A and 5C showed that high glucose could promote the tube formation, and the formation was inhibited by ASK1 and p38 inhibitor. Meanwhile, the tube meshes, nodes and tube length was counted. The result showed that HRMECs in high glucose could form more meshes and nudes than control; meanwhile, the tube total length in HG group was higher than control group (Figure 5B and D). However, after blocking ASK1 and p38, the tube meshes, nudes and tube length was reduced comparing with HG group (Figure 5B and D). These data suggested that ASK1/p38 signal axis participated in tube formation of retinal microvascular endothelial.

**Figure 5.**
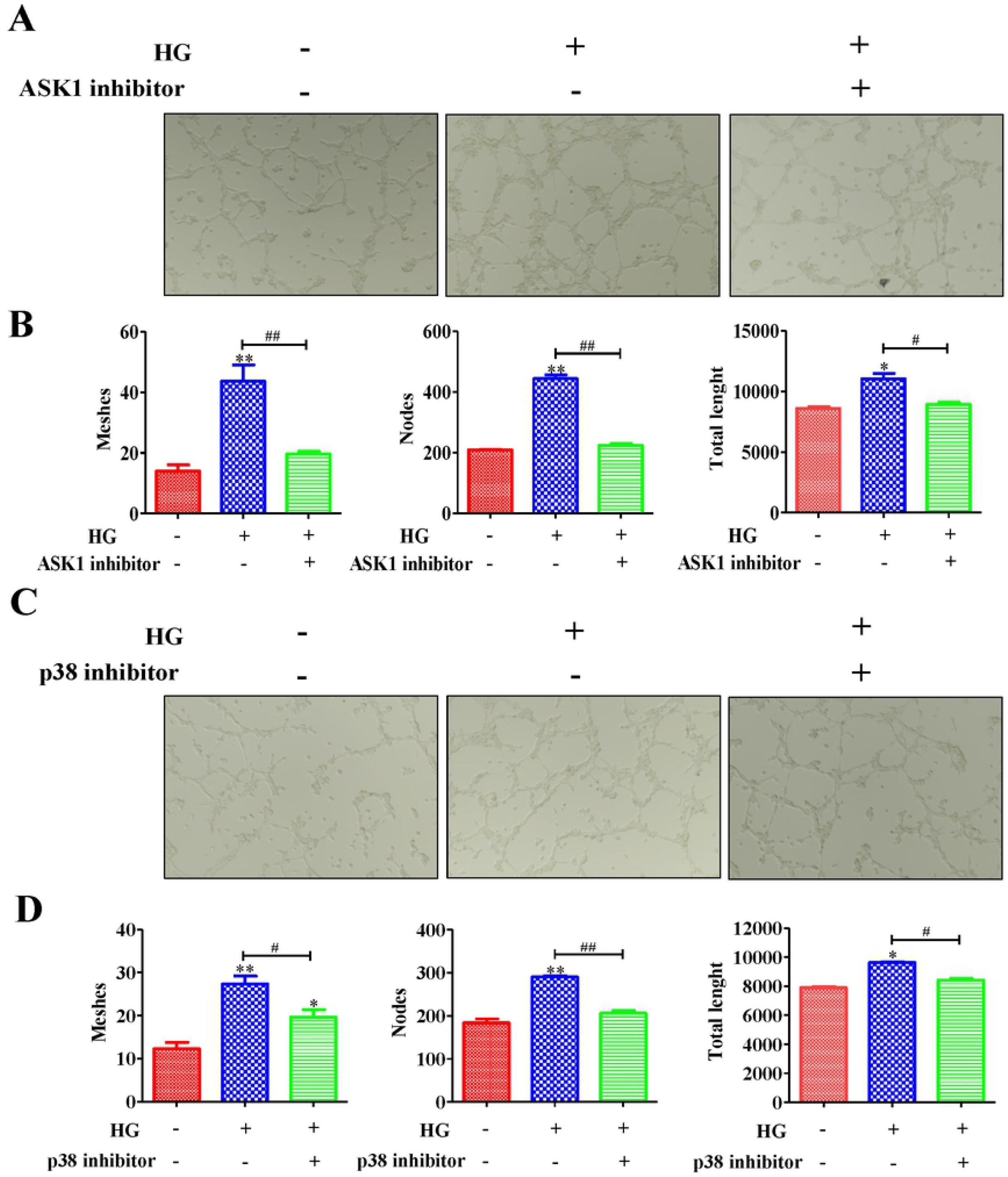
ASK1/p38 signal axis regulated the tube formation of retinal microvascular endothelial. (A) The tube formation was obviously enhanced in HG HRMECs and was inhibited by blocking ASK1. Then, the (B) tube meshes, nodes and tube length was counted, and the results exhibited that HRMECs in high glucose could form more meshes and nudes than control; meanwhile, the tube total length in HG group was higher than control group. However, after blocking NLRP3, the tube meshes, nudes and tube length was reduced comparing with HG group. In addition, (C) The tube formation was inhibited by blocking p38 and the (D) tube meshes, nodes and tube length was counted; the result showed that blocking p38 could inhibit the tube formation. *P < 0.05 and **P < 0.01 compared with the control. #P < 0.05 and ##P < 0.01 compared with the relative HG-induced HRMEC cell group.

### The angiogenesis was inhibited through blocking ASK1 and p38

Next, we investigated whether ASK1 and p38 mediated the angiogenesis. The results showed that angiogenesis related marker IB4 was obviously increased in HD model (Figure 6A and B); However, IB4 expression was inhibited through blocking ASK1 and p38 (Figure 6A and B). This data suggested that ASK1 and p38 participated in angiogenesis in DR.

**Figure 6.**
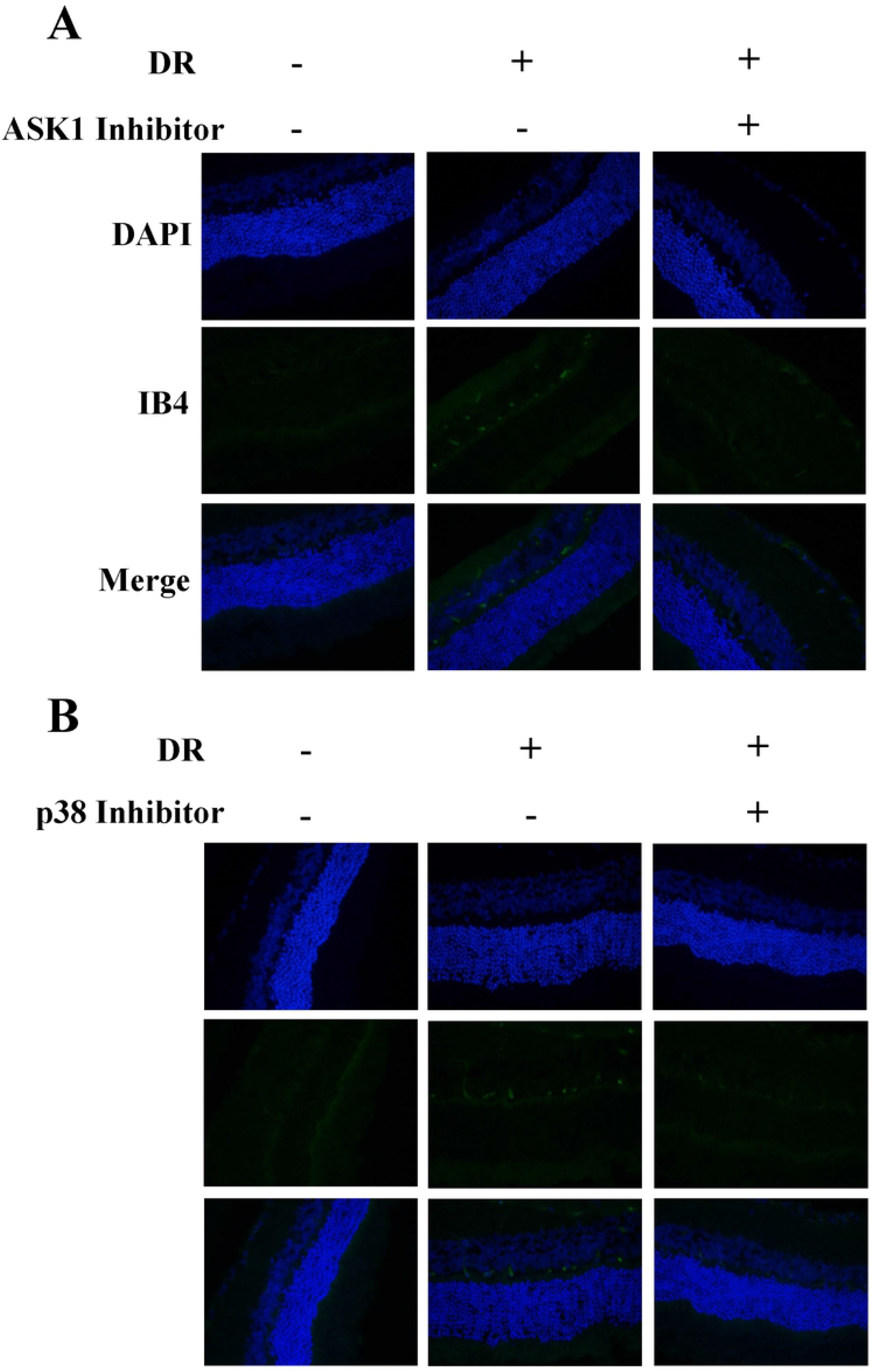
The angiogenesis was inhibited through blocking ASK1 and p38. Angiogenesis related marker IB4 was obviously increased in HD model; However, IB4 expression was inhibited through blocking (A) ASK1 and (B) p38.

## Discussions

DR is a leading cause of blindness in working-age population all the world. It is induced by diabetes, which causes the pathology of retinal capillaries, arterioles and venules, and the subsequent effects of leakage from or occlusion of small vessels [22–24]. Laser therapy is the mainly effective therapy for preservation of sight in proliferative retinopathy [24]; however, it is poor to reverse visual loss. Hence, it is still to find the new therapies to improve the effective of DR.

In this study, we had uncovered the NLRP3 played an important part in aberrant retinal angiogenesis in diabetic retinopathy. We firstly found that ASK1/p38-mediated NLRP3 inflammasome signaling pathway contributed to aberrant retinal angiogenesis in diabetic retinopathy. The inflammatory related cytokines (IL-6, TNF-α, IL-1β) mRNA expression levels were up-regulated in DR (Figure 1A); meanwhile, vascular related marker CD31 and cytokine VEGF were also increased in DR (Figure 1B-C). After blocking NLRP3, ASK1 or p38, the expression levels of inflammatory cytokine was down-regulated and vascular related marked IB4 expression was decreased.

It is the crucial reason that heperglycemia lead to a series of inflammatory mediator in diabetes, which can further parainflammatory response and finally damage retinal microvascular [25]. Chronic inflammation is one of the key that triggers in the pathogenesis of DR, and the inflammation response reduction can alleviate the development and progression of DR [26]. Related reports show that NLRP3 plays a crucial key in the development of chronic inflammatory response through secreting related cytokines, such as IL-6, TNF-α, IL-1β [27]. The activation of NLRP3 contributes to all kinds of development and progression in chronic inflammatory disease [28]. In this study, our results showed that NLRP3-mediated tissue inflammatory response promoted microvascular proliferation in retina. In DR animal model, the protein expression of NLRP3 was up-regulated in DR group (Figure 2A). Then, the DR group was treated with NLRP3 inhibitor; after inhibitor treatment, the inflammation cytokines IL-6, TNF-α and IL-1β mRNA expression levels were decreased comparing with DR group without treated with inhibitor (Figure 2B). Meanwhile, CD31 expression level was reduced in inhibitor treatment group (Figure 2C). Moreover, the VEGF secretion level was reduced after administrated with NLRP3 inhibitor (Figure 2D). Moreover, the NLRP3 protein expression level in HG HRMECs group was up-regulated, and inhibited after treated inhibitor (Figure 2E). In HG HRMECs groups, IL-6, TNF-α and IL-1β mRNA expression levels were enhanced comparing with control, and inhibited by NLRP3 inhibitor (Figure 2F). The result of VEGF secretion levels in cells was similar with DR animal model (Figure 2G).

ASK1 is an apoptosis-related protein, which is activated in response to a variety of stress-related stimuli *via* distinct mechanisms and activates MKK4 and MKK3, which in turn activate JNK and p38 [29]. Many reports illustrate that ASK1 can contribute to the development and progression of inflammatory response [30,31]. For example, the bacterial infection-engaged inhibition of ASK1 is responsible for regulating Erk1/2- and p38-MAPKs activation, but not JNK-MAPK signaling [31,32]. This previous studies suggest that ASK1 and p38 have close relationship in inflammatory response. In addition, some drugs can protect the retinal photoreceptor cells through activating p-Erk1/2/Nrf2/Trx/ASK1 signaling pathway in diabetic mice [33]. This report suggested that ASK1 play an important role in diabetic related disease [33]. In this study, our data showed that the expression level of inflammatory some NLRP3 was up-regulated through activating ASK1/p38 signal axis; meanwhile, ASK1/p38 signal axis contribute to tube formation of retinal microvascular endothelial and development of angiogenesis in DR. We found that ASK1 and p38 protein expression levels were up-regulated in DR (Figure 4A-B); however, blocking ASK1 and p38, the NLRP3 and related cytokine (IL-6, TNF-α, IL-1β) protein expression levels were down-regulated (Figure 4C-D). In addition, the tube formation of retinal microvascular endothelial was inhibited through blocking ASK1 and p38. All of this data showed that ASK1 and p38 mediated the NLRP3 inflammasome signaling pathway contributed to aberrant retinal angiogenesis in diabetic retinopathy.

## Conclusions

In conclusion, we found that DR could cause the inflammatory response and microvascular proliferation. NLRP3 contributed to DR-mediated inflammatory development and progression, which promoted the inflammatory related cytokine expression. Meanwhile, it could promote the tube formation of retinal microvascular endothelial and angiogenesis. Moreover, further research showed that NLRP3 mediated aberrant retinal angiogenesis in diabetic retinopathy was regulated by ASK1 and p38.

## Data Availability Statement

All relevant data are within the manuscript.

## Funding

This research was financially supported by National Natural Science Foundation of China (81700852), the Young Talent’s Subsidy Project in Science and Education of the Department of Public Health of Jiangsu Province (QNRC2016140) and Funded Project of the Wuxi Municipal Health and Family Planning Commission (Q201623)

